# RegCyanoDB: a database of regulatory interactions in cyanobacteria

**DOI:** 10.1101/117127

**Authors:** Ajay Nair, Madhu Chetty, Nguyen Xuan Vinh

**Affiliations:** IITB-Monash Research Academy, Indian Institute of Technology Bombay, Mumbai, 400 076, India; Faculty of Information Technology, Monash University, Melbourne, VIC 3800, Australia; Chemical Engineering Department, Indian Institute of Technology Bombay, Mumbai, 400 076, India; Faculty of Science and Technology, Federation University, VIC 3841, Australia; School of Computing and Information Systems, The University of Melbourne, VIC 3010, Australia

**Keywords:** Cyanobacteria, *Escherichia coli*, gene regulatory network, transcriptional regulation, transcription factors, regulatory interactions, regulatory interaction mapping, Bioinformatics

## Abstract

**Background:** Cyanobacteria are photoautotrophic organisms with environmental, evolutionary, and industrial importance. Knowledge of its regulatory interactions are important to predict, optimise, and engineer their characteristics. However, at present, very few of their regulatory interactions are known. The regulatory interactions are known only for a few model organisms such as *Escherichia coli* due to technical and economical constraints, which are unlikely to change soon. Thus, mapping of regulatory interactions from well-studied organisms to less-studied organisms by using computational techniques is widely used. Reverse Best Hit (RBH), with appropriate algorithm parameters, is a simple and efficient method for detecting functional homologs.

**Description:** We predict the regulatory interactions in 30 strains of cyanobacteria using the known regulatory interactions from the best-studied organism, *E. coli.* RBH method with appropriate parameters is used to identify the functional homologs. An interaction is mapped to a cyanobacterial strain if functional homologs exist for a known transcription factor and its target gene. The confidence of the detected homologs and interactions are also provided. Since RBH is a conservative method, homolog-grouping is performed to recover lost putative interactions. A database of the predicted interactions from all the 30 strains of cyanobacteria is constructed.

**Conclusion:** RegcyanoDB contains 20,280 interactions with confidence levels for 30 cyanobacterial strains. The predicted regulatory interactions exhibit a scale free network topology as observed in model organisms. The interacting genes in *E. coli* and cyanobacteria are mostly found to have the same gene annotation. This database can be used for posing novel hypotheses and validation studies in wet-lab and computational domains.

The database is available at http://www.che.iitb.ac.in/grn/RegCyanoDB/

## 1 Introduction

Cyanobacteria, or the blue-green bacteria, are photoautotrophic organisms credited with changing the Earth's atmosphere to oxygen rich condition. It is believed that the photosynthetic machinery in plants and algae, the chloroplast, has evolved from cyanobacteria by endosymbiosis. Cyanobacteria are the model organism for studying photosynthesis, as well as nitrogen and carbon assimilation. They are believed to play an important role in marine nitrogen-fixing cycle [1,2]. Presently, there is enormous interest in using cyanobacteria for biofuel and hydrogen production [3-5]. The algal species and cyanobacteria have the highest biofuel production per unit area [3] and have higher growth and photosynthetic rates [5]. *N*_2_-fixing cyanobacterial strains have simple growth conditions and have the simplicity of prokaryotic genome compared to eukaryotic algae [5]. Thus, they are a promising sustainable alternative to our energy requirements.

For biofuel applications, understanding the metabolic and genetic factors involved in maximising the productivity in an organism are of great interest [3,6]. Organisms respond to varying environmental conditions by regulating the cellular protein production. In prokaryotes, the regulation of the proteins is largely carried out by the transcriptional networks. Thus, understanding the transcriptional regulatory network is crucial in optimising the culture conditions and for metabolic or genetic manipulation.

Although around 2000 completely sequenced genomes are reported in GOLD database [7], their functional characterization and regulatory interactions are lagging behind. Even for the model organism *Escherichia coli* K-12, whose regulatory network is best understood of all the living organisms [8], only about one-third of the genes have experimentally validated interactions (RegulonDB database, June 2012). The main reason for this limited information is that there are many technical and organizational issues associated with finding regulatory interactions experimentally [9]. Further, compared to metabolic networks, using comparative genomics in regulatory networks is more challenging as they are less conserved, very plastic, and the transcription factors evolve fast [10–13]. As a result, our current knowledge of the regulatory networks in prokaryotes is limited to only a few model organisms. These are available in public databases such as RegulonDB [8] and EcoCyc [14] for *E. coli*; DBTBS [15] for *Bacillus subtilis*; MtbRegList [16] and MycoRegNet [17] for *Mycobacterium tuberculosis*; and CoryneRegNet [18] for corynebacteria. RegTransBase [19] contains manually-curated, experimentally-verified interactions for 128 microbes, while PRODORIC [20] contains the regulatory information of many prokaryotes but mainly *E. coli, B. subtilis,* and *Pseudomonas aeruginosa.* Due to difficulties involved in obtaining regulatory interactions experimentally, mapping regulatory interactions from model organisms to others using computational techniques is widely used [9,10,16–18,21–23]. While generic databases like RegTransBase [19] and PRODORIC [20] contain interactions for cyanobacteria, these interactions are very few. Hence, it has become imperative to develop a database which contains computationally predicted regulatory interactions for this organism.

RegCyanoDB is thus, the first database of regulatory interactions in cyanobacteria. The regulatory interactions are mapped using *E. coli* as the reference organism. This database also provides the confidence level of the predicted interactions based on the quality of the sequence alignment.

## 2 Computational methods for regulatory interaction mapping

Several studies have characterized the two main assumptions in computational transfer of regulatory interactions: (i) the function of a new protein can be predicted using its sequence similarity to a known protein; and (ii) for a known transcription factor(TF) and its target gene(TG) in the ‘source’ organism, the interaction is conserved in the ‘target’ organism if there exist functional homologs for both the TF and its corresponding TG. The concept of “interologs”, the orthologous pair of interacting proteins, was reported [24] for transferring protein-protein interactions between organisms. This concept was extended to a large scale study [23] that introduced the concept of “regulog”, the conserved protein-DNA interactions in different organisms. It was shown that if a TF has a homolog in another organism with 30-60% or better sequence identity, the binding site of the homolog is conserved and for identities above 80%, all the protein-protein interactions are conserved. The sequence identity values reported in [23] also matched observations in [25], which noted that the pairwise alignment of two sequences correlated the structural alignment when the sequence identity is above 25-30%.

Benchmark studies [26–28] and reviews [29, 30] have analysed the various ortholog detection methods. Recent benchmark studies have shown that reverse best hit (RBH) is as good or even superior to other methods [26,27]. Note that RBH is also referred to as reciprocal best hit, or bidirectional best hit (BBH), or symmetrical best hit (SymBeT). RBH is a pairwise sequence alignment method that uses the concept of orthologous genes which is operationally defined as the gene pair having the best sequence similarity between all the genes in two genomes [13,31]. Thus, if a protein *P*1_*x*_ in the first organism picks protein *P*2_*y*_ as its best hit in a sequence similarity search against all the proteins in the second organism, and if *P*2_*y*_ picks *P*1_*x*_ as its best hit among all the proteins in the first organism, then *P*1_*x*_ and *P*2_*y*_ are called the RBH of each other. It is important to note that the sequence similarity between the two proteins should have sufficient statistical significance [31]. Among the different methods for ortholog detection, sequence similarity based methods like RBH are strong predictors of functional relatedness [26] and appropriate choice of algorithm parameters yield good results [29,32,33]. Orthologs are generally accepted to be functionally equivalent [13,30,31].

Difficulties for functional prediction using RBH arise when there is domain shuffling, presence or absence of domains, gene duplication, gene loss, and horizontal gene transfer [25,30]. The possible error due to changes in protein domains are addressed by considering the coverage of the pairwise alignment. Different implementations have used different coverage criteria, such as, 80% coverage [23,34], or 70% coverage along with protein domain information [12], or 60% coverage with 60% identity in the alignment region [10], or 50% coverage with relevant e-value cut-off [33].

Many-to-one or one-to-many relations caused by gene deletions and duplications cannot be considered in RBH method and low sequence similarity of alignment will require additional methods like conserved gene neighbourhood analysis [29,30,35]. Since RBH considers only the best hit in both the directions, it is considered as a conservative method. Therefore, clustering or grouping of homologous proteins is used to recover the false negatives [10,35]. Additional constraints, like minimum sequence identity and coverage are considered to minimize false positives [10,12,23,25,33].

BLAST [36, 37] is by far the most popular method for sequence similarity searches. Ortholog detection using BLAST with different parameter cut-offs such as e-value, raw-score, bit-score, and identity give different but essentially overlapping results [29]. The cut-off for sequence similarity should be statistically significant but not too stringent [31]. Smith-Waterman implementation with e-value cut-off was found to be a good measure of structural similarity between the proteins [32] and Smith-Waterman alignment with soft-filtering in BLAST was reported to give best results for RBH [33].

These results form the basis for parameter selection for the algorithms and for predicting the confidence of regulatory interactions in this work.

## 3 Construction and content

The regulatory interactions in 30 strains of cyanobacteria were mapped from the known regulatory network in *E. coli* K-12. The database construction procedure for a single cyanobacterial strain is shown in figure 1. The experimentally characterised regulatory interactions in *E. coli* were obtained from RegulonDB [8] which had 3920 interactions, 183 TFs, and 1563 unique mRNA genes (as on June 2012). Interactions with other gene products like tRNA, rRNA, and ncRNA were ignored. The genes reported in the interactions were identified mainly using RSAT [38,39] and the rest from NCBI RefSeq [40] and UniProtKB [41]. Protein sequences of *E. coli* were obtained mainly from NCBI RefSeq and the rest from UniProtKB. Complete protein sequences of the 30 cyanobacterial strains were obtained from NCBI RefSeq. The list of strains in Table 1, include all the major orders such as Chroococcales (21 strains), Nostocales (4 strains), Prochlorales (2 strains), Gloeobacteria (1 strain), and Oscillatoriales (1 strain).

**Figure 1:**
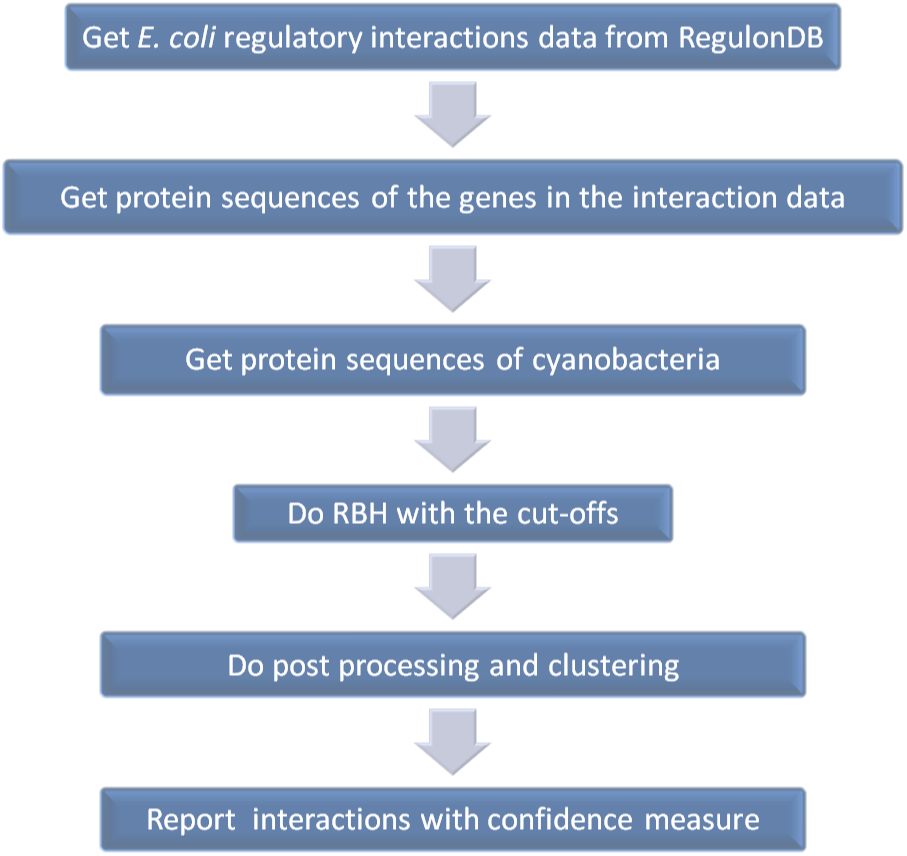
Procedure for obtaining regulatory interactions for a cyanobacterial strain

**Table 1:** The 30 strains of cyanobacteria used in the study.

| Order | Strain |
| --- | --- |
| Chroococcales | *Cyanothece* sp. ATCC 51142<br>*Cyanothece* sp. PCC 7424<br>*Cyanothece* sp. PCC 7425<br>*Cyanothece* sp. PCC 7822<br>*Cyanothece* sp. PCC 8801<br>*Cyanothece* sp. PCC 8802<br>*Microcystis aeruginosa* NIES-843<br>*Synechococcus elongatus* PCC 6301<br>*Synechococcus* sp. CC9311<br>*Synechococcus* sp. CC9605<br>*Synechococcus* sp. CC9902<br>*Synechococcus* sp. JA-3-3Ab<br>*Synechococcus* sp. PCC 7002<br>*Synechococcus* sp. RCC307<br>*Synechococcus* sp. WH 7803<br>*Synechococcus* sp. WH8102<br>*Synechocystis* sp. PCC 6803 substr. PCC-N<br>*Synechocystis* sp. PCC 6803 substr. PCC-P<br>*Synechocystis* sp. PCC 6803 substr. GT-I<br>*Thermosynechococcus elongatus* BP-1<br>*Cyanobacterium* UCYN-A |
| Gloeobacteria | *Gloeobacter violaceus* PCC 7421 |
| Nostocales | *Anabaena* sp. PCC 7120/Nostoc sp. PCC7120<br>*Anabaena variabilis* ATCC 29413<br>*Nostoc azollae* 0708<br>*Nostoc punctiforme* ATCC 29133 |
| Oscillatoriales | *Trichodesmium erythraeum* IMS101 |
| Prochlorales | *Prochlorococcus marinus* str. MIT 9215<br>*Prochlorococcus marinus* SS120 |

RBH was implemented in standalone BLAST (version 2.2.26+) from NCBI with the following parameters. The e-value cut-off 1e-04 was decided by the database size [32] of ~6000 protein sequences for a single BLAST run. Smith-Waterman alignment and soft-filtering were enabled for optimal alignment results [32,33].

After obtaining the protein sequences, BLAST of *E. coli* proteins against proteins of a selected strain of cyanobacteria and the reverse BLAST of the cyanobacterial proteins against the *E. coli* proteins were performed. The RBH proteins were selected from the BLAST results and graded for functional homolog confidence. The classification of confidence level is shown in Table 2. For a pairwise sequence alignment, if the identity was at least 80% and alignment length covered at least 80% of the two proteins, the homolog confidence was reported ‘high’. As discussed earlier, at this sequence identity level the protein-protein interactions and protein-DNA interactions are expected to be conserved [23]. If the identity is between 60 – 80% and coverage is at least 80%, the homolog confidence was assigned ‘good’, as protein-DNA interactions are expected to be conserved and many protein-protein interactions are also conserved [23]. Similar cut-off had been reported to give optimal results for regulatory interaction mapping [10]. Homolog confidence was reported as ‘moderate’ for identity between 25 – 60% and coverage greater than 60%. At this identity levels, protein-DNA interactions could be conserved [23] and sequence alignment correlates structural alignment [25]. All other RBH cases were reported as ‘low’ confidence homologs as the confidence in their function cannot be decided by quality of sequence alignment alone.

**Table 2:** Confidence levels for a functional homolog using RBH. The intervals are based on the pairwise alignment parameters of sequence identity (SI)% and the coverage of the alignment or the alignment length (AL) in % for both proteins from an RBH.

| Confidence level | Parameters |
| --- | --- |
| High | SI > 80% |
|  | AL > 80% |
| Good | SI > 60% |
|  | AL > 80% |
| Moderate | SI > 25% |
|  | AL > 60% |
| Low | Other RBH |

For the proteins in *E. coli* which did not have a RBH, ‘homolog-groups’ were created. In this procedure, homolog confidence was computed for all the hits of an *E. coli* protein *PE_x_* against the selected cyanobacteria. All the homolog proteins in the hit having confidence of ‘moderate’ or better were listed together in *‘Cluster_x_*’ as the putative functional homologs of *PE_x_*. The confidence level of the homolog-group was assigned as the highest homolog confidence shown by any member in the group. This procedure is similar in concept to the reported clustering methods [10, 35] and is described as follows. RBH cannot extract the functional homologs in the presence of gene deletions and duplications. However, proteins with sufficient identities and coverage in sequence alignment can be functionally equivalent. So, when RBH is not present, all the homolog proteins that show sufficient identity and coverage in pairwise sequence alignment are reported as putative functional homologs. Thus, false negatives from the conservative RBH method can be recovered. However, since there are chances of false positives also being present, the results with homolog-group procedure are reported separately.

After identifying the functional homologs using RBH and homolog-grouping, the TF-TG interactions were obtained using the interactions reported in RegulonDB database. For a TF-TG interaction reported in RegulonDB, corresponding interaction is considered as conserved in the selected cyanobacteria, if a homolog or a homolog-group can be identified for both the TF and TG in the cyanobacteria. The confidence of the interaction is assigned as the lowest of the homolog confidence level between the homolog TF and the TG. For example, if the TF homolog confidence was ‘moderate’ and the TG homolog confidence was ‘high’, the interaction confidence is taken as ‘moderate’. Wherever possible, the homolog proteins identified in cyanobacteria were also verified by their annotations to their *E. coli* protein annotations. The results obtained from RBH and obtained by both RBH and homolog-grouping are reported separately in database. This procedure was repeated for all the other strains of cyanobacteria to create complete database.

For illustration, the predicted regulatory interactions of *Cyanothece* sp. ATCC 51142 and *Acaryochloris marina* MBIC11017, visualized using Cytoscape [42], are shown in figures 2 and 3. It can be seen that the network approximates a scale-free topology in which a small number of TFs regulate a large number of TGs, forming network hubs, and a large number of TFs control only a small number of TGs. Further, a large number of TGs are regulated by only a small number of TFs and a small number of TGs are controlled by many TFs. Other cyanobacterial strains also showed similar network topology. This is similar to the gene regulatory network topology reported for other well studied organisms [43,44].

**Figure 2:**
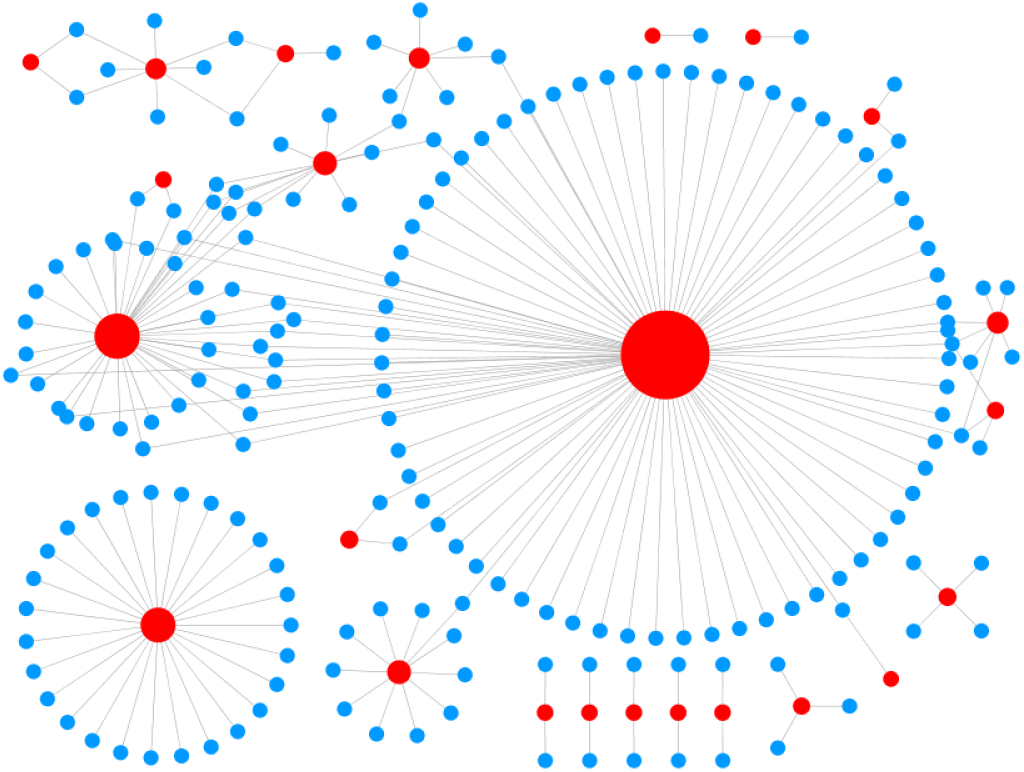
Regulatory network of *Cyanothece* sp. ATCC 51142. The red-nodes represent transcription factors in which the node-size is proportional to the out-degree. The blue-nodes represent target genes and edges represent the interactions. The network topology shows a scale free structure.

**Figure 3:**
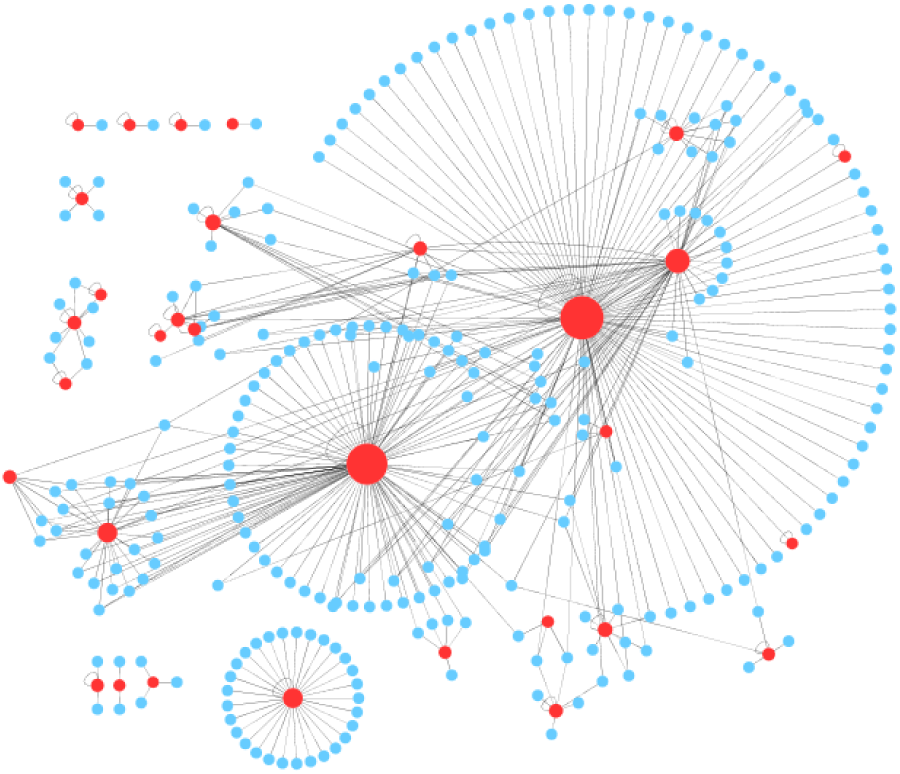
Regulatory network of *A. marina* MBIC11017. The red-nodes represent transcription factors whose node-size is proportional to the out-degree. The blue-nodes represent target genes and edges represent the interactions. The network topology shows a scale free structure.

Gene regulatory interactions in different strains of cyanobacteria predicted using RBH method and a combination of RBH and homolog-grouping method are shown in figure 4. The absolute number of interactions predicted for the cyanobacterial strains are shown in figure 4(A). The relative number of interactions predicted for the different strains are shown in figure 4(B) which can be used for comparison of the conservation of interactions among the different strains. It is the ratio of the number of interactions predicted in a strain to the average number of interactions obtained for all the strains, in each method. It can be observed that the strains of Prochlorales and *Cyanobacterium* UCYN-A, which have the smallest genomes among all strains, have the lowest number of the detected interactions. *Nostoc azollae* 0708, which is a symbiont to fern *Azolla*, has the lowest number of interactions among the Nostocales strains. This probably represents the gene loss during symbiosis [45]. Interactions are relatively fewer in Synechococcus strains.

**Figure 4:**
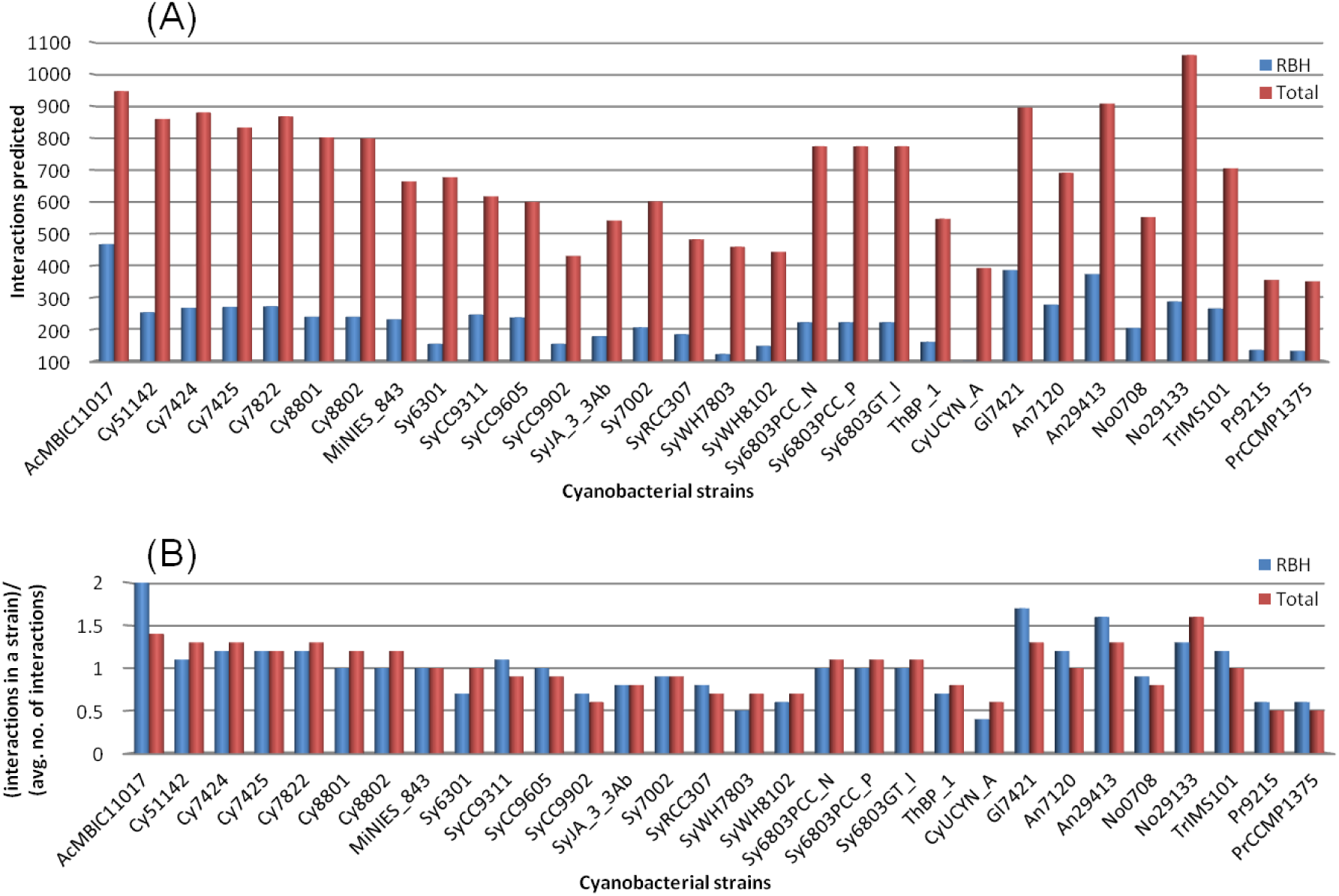
Regulatory interactions predicted in all cyanobacterial strains. All the 30 strains of cyanobacteria are shown in horizontal-axis. Red bars represent the interactions for RBH method and blue bars show total results for both RBH and homolog-grouping methods. (A) The absolute number of interactions predicted for all the 30 strains, using both the methods, are shown. (B) The relative number of interactions predicted in the different strains; vertical-axis is the ratio of interactions in a strain to the average number of interactions obtained for all the strains in each method.

## 4 Utility and Discussion

### 4.1 RegCyanoDB user interface

The regulatory interactions in different cyanobacterial strains are available for download from the database website. Selecting ‘Downloads’ from the main page, and then choosing a specific strain (such as *Cyanothece* sp. ATCC 51142), display links to text files containing regulatory interactions. These files are formatted to aid manual or computational analysis. The ‘Downloads’ page for *Cyanothece* sp. ATCC 51142 is shown in figure 5.

**Figure 5:**
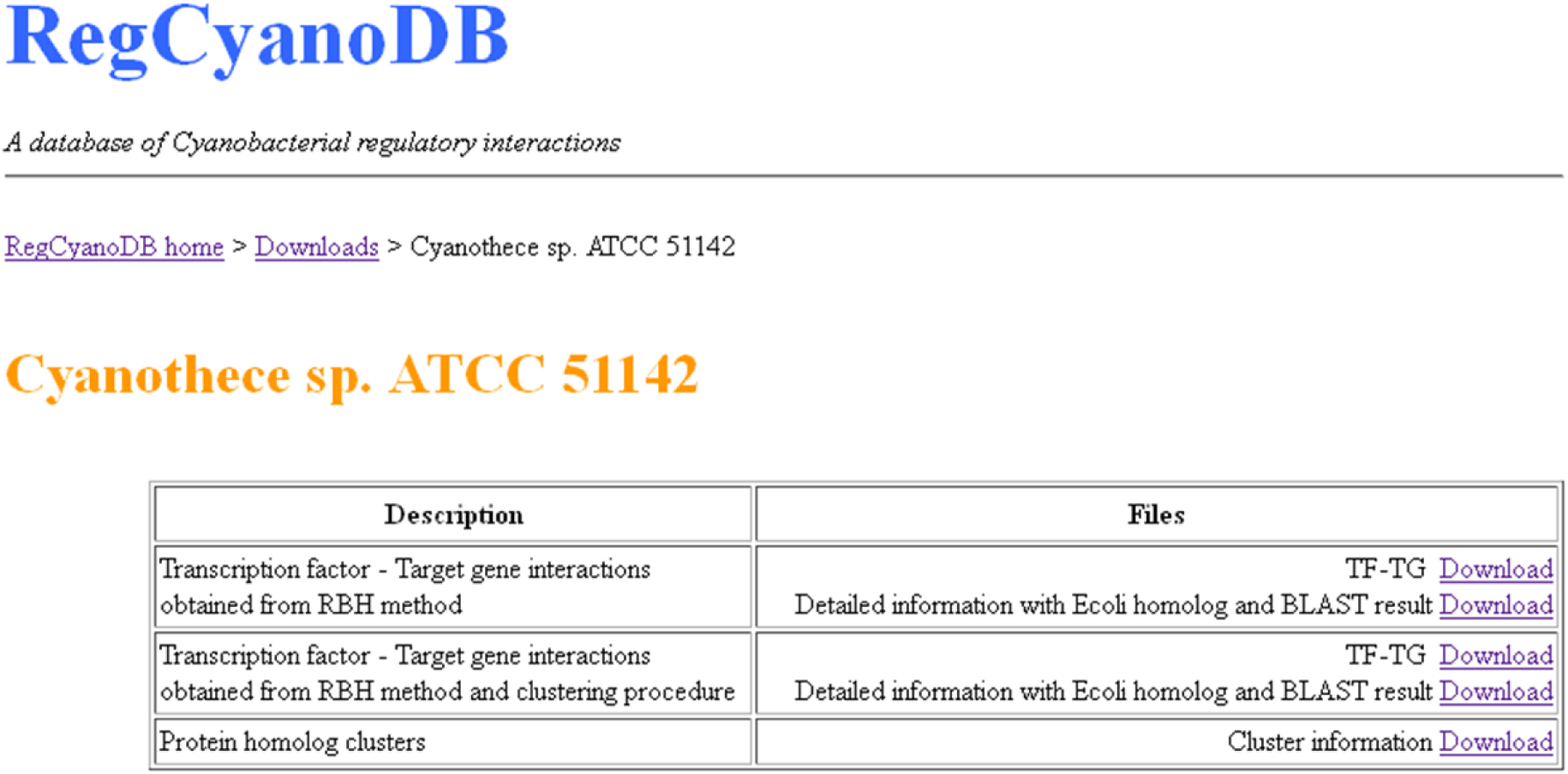
‘Downloads’ section for *Cyanothece* sp. ATCC 51142 which shows the regulatory interaction information for cyanobacterial strain *Cyanothece* sp. ATCC 51142 available at the website.

The ‘Transcription factor – Target gene interactions’ section gives the RBH interactions alone, with no information from the homolog-grouping procedure. This is expected to contain only conservative number of interactions. ‘TF-TG’ section reports just the reference number of TFs and TGs and their interaction confidence. The ‘Detailed information’ section gives the protein name, the *E. coli* protein, and the BLAST results.

The interactions detected using the homolog-grouping procedure along with the RBH method is in second row. This section again contains two files, one concise and other detailed, as describe previously.

The details of the proteins present in the homolog-groups and their ‘Cluster numbers’ are given in the ‘Protein homolog-groups’ section.

### 4.2 RegCyanoDB significance

To the best of our knowledge, RegCyanoDB is the first dedicated regulatory interaction database for cyanobacteria. While there are other databases for cyanobacteria that give information about the transcription factor families [22], protein-protein interactions in specific strains [46], operons [47], and genome details [48], none of these provide regulatory interaction information. Currently, only a few hundred regulatory interactions for cyanobacteria are available in the public databases like PRODORIC and RegTransBase.

This database predicts 20, 280 interactions for the 30 strains of cyanobacteria along with confidence levels in the predicted homologs and interactions.

The regulatory interactions of *E. coli* in RegulonDB database had been used for upto 100 different applications [49], both experimental and computational. Since the cyanobacteria database reported here is computationally predicted, it can serve as the first step for targeted wet-lab experiments studying the regulatory interactions in this organism. Since the microbes predominantly use transcriptional regulation to adapt itself, the information in the website can also be used to generate new hypothesis about the characteristics and phenotype of cyanobacteria [50].

The information from the database will also be useful to understand and analyse the microarray gene expression data, predict upstream binding locations of different TFs, detect the TFBS motifs, analyse protein expression, and study regulatory interactions in different strains of cyanobacteria. Other important investigations in bioinformatic applications such as assigning protein function, uncovering novel interactions, and studying operons [47,51,52], will be aided by this database. The structure of the gene regulatory network in cyanobacteria at different levels, e.g. individual interactions, network motifs, and also at the global level can be studied. Current gene regulatory network modelling techniques which are limited due to ‘curse of dimensionality’, i.e. too many variables and too few genes, will also be benefited as these known interactions and the transcription factors can be used as a-priori knowledge in the modelling process.

## 5 Conclusions

RegCyanoDB provides the computationally predicted regulatory interactions in cyanobacteria, mapped from the most well-studied organism, *E. coli*. A total of 20, 280 interactions with confidence levels, have been reported for the 30 strains of cyanobacteria. These confidence levels will give a better idea for using the interactions in experimental or computational applications. Further, we observe that the regulatory interactions obtained in the cyanobacterial strains approximate a global scale-free network topology as reported for other model organisms.

## Competing interests

The authors declare that they have no competing interests.

## Author contributions

AN and MC conceptualised the work. AN collected the data, developed the algorithms, carried out the experiments and developed the database. NXV provided critical suggestions. AN, NXV and MC drafted the manuscript. All authors read and approved the final manuscript.

## Acknowledgements

AN performed part of this work in Pramod Wangikar's lab. He thanks Pramod Wangikar for taking part in conceptualisation and some of the discussions on this work. He thanks Dilip Durai for feedback on Perl codes and S Krishnakumar for general discussions.

